# Odd skipped-related 1 (Osr1) identifies muscle-interstitial fibro-adipogenic progenitors (FAPs) activated by acute injury

**DOI:** 10.1101/249003

**Authors:** Jürgen Stumm, Pedro Vallecíllo Garcia, Sophie vom Hofe-Schneider, David Ollitrault, Heinrich Schrewe, Aris N. Economides, Giovanna Marazzi, David A. Sassoon, Sigmar Stricker

## Abstract

Fibro-adipogenic progenitors (FAPs) are resident mesenchymal progenitors in adult skeletal muscle that support muscle repair, but also give rise to fibrous and adipose infiltration in response to disease and chronic injury. FAPs are currently identified using cell surface markers that do not distinguish between quiescent FAPs and FAPs actively engaged in the regenerative process. We have shown previously that FAPs are derived from cells that express the transcription factor Osr1 during development. Here we show that adult FAPs express Osr1 at low levels and frequency, however upon acute injury FAPs reactivate Osr1 expression in the injured tissue. Osr1^+^ FAPs are enriched in proliferating and apoptotic cells demonstrating that Osr1 identifies activated FAPs. *In vivo* genetic lineage tracing shows that Osr1^+^ activated FAPs return to the resident FAP pool after regeneration as well as contribute to adipocytes after glycerol-induced fatty degeneration. In conclusion, reporter LacZ or eGFP-CreERt2 expression from the endogenous Osr1 locus serves as marker for FACS isolation and tamoxifen-induced manipulation of activated FAPs.

**Summary statement:** Expression of Osr1 specifically in muscle interstitial fibro-adipogenic progenitors (FAPs) activated by acute injury provides a tool to isolate and trace this population.

## Introduction

The remarkable regenerative potential of skeletal muscle relies on myogenic stem cells (satellite cells), however other interstitial populations play a critical supportive role (Bentzinger et al., 2013; Pannerec et al., 2012; Uezumi et al., 2014b). Amongst these, fibro-adipogenic progenitors (FAPs) have attracted immense attention in the past years. FAPs are muscle-interstitial resident mesenchymal progenitor cells that have the capacity to provide a pro-myogenic environment for muscle regeneration (Joe et al., 2010) but also contribute directly to fibrotic degeneration and fatty infiltration in diseased or degenerating muscle (Lemos et al., 2015; Uezumi et al., 2010; Uezumi et al., 2011). As such, FAPs are important cell targets for therapeutic approaches (Contreras et al., 2016; Gonzalez et al., 2017; Lemos et al., 2015; Mozzetta et al., 2013). FAPs are activated upon injury to proliferate (Joe et al., 2010; Uezumi et al., 2010) and are cleared by apoptosis in the course of regeneration (Lemos et al., 2015). The intrinsic mechanisms of activation and pro-myogenic function as well as the mechanisms that promote fibrotic or adipogénie conversion are not well understood. Murine FAPs were characterized using different cell surface marker combinations. Joe et al. (2010) used the combination of lin^−^;Scal^+^;CD34^+^ or equivalently lin^−^;α7-integrin^−^;Scal^+^ to isolate FAPs, while Uezumi et al (2014a; 2010) used lin^−^;PDGFRα^+^ to isolate FAPs from mouse and human muscle. Both, the Sca1^+^ and PDGFRα^+^ populations appear to largely overlap (Uezumi, Ikemoto-Uezumi and Tsuchida, 2014b). In addition, FAPs show overlap with Tcf4^+^ cells originally defined as muscle connective tissue fibroblasts (Murphy et al., 2011; Vallecíllo-Garda et al., 2017) as well as with the PDGFRα^+^ subpopulation of PICs (PW1^+^ interstitial cells). PICs are Sca1^+^ and CD34^+^ and are marked by expression of the paternally imprinted gene PW1 (Peg3), which is a general stem cell / progenitor cell marker (Berg et al., 2011; Besson et al., 2011). PICs were originally characterized as an interstitial multipotent population distinct from satellite cells (Mitchell et al., 2010). Later it was shown that PICs can be divided into PDGFRα^−^ myogenic PICs and PDGFRα^+^ adipogenic PICs that completely overlap with FAPs (Pannerec et al., 2013). The above mentioned markers label tissue-resident quiescent FAPs as well as FAPs activated upon injury or disease. To date, no molecular marker has been found to identify injury-activated FAPs, and no tool exists to specifically purify or manipulate this population, precluding analyses as well as genetic manipulation or *in vivo* lineage tracing of injury-activated FAPs. The identification of an activated FAP-specific molecular marker promises to greatly facilitate our understanding of the intrinsic mechanisms of FAP activation, function, and differentiation under normal or pathological conditions. Here we show that FAPs become positive for the transcription factor Osr1 (Odd skipped-related 1) in response to injury and that Osr1 expression can be used to follow, isolate, and genetically mark activated FAPs during the pathological and as normal muscle repair.

## Results and Discussion

### Osr1 is expressed in a small number of adult FAPs

During development Osr1 marks a lateral plate mesoderm-derived population of fibro-adipogenic cells that is also a source for adult Sca1^+^ and PDGFRα^+^ FAPs (Vallecillo-García et al., 2017). However, *Osr1* expression declines during development and early postnatal life in mice, and eGFP expressed from the Osr1 locus (Osr1^GCE^ mouse line; Mugford et al., 2008) is only detectable *in situ* by antibody staining during early postnatal life but is below detectable levels in adults (Vallecillo-García et al., 2017). To increase detection sensitivity, we inserted a β-galactosidase (β-Gal) reporter into the *Osr1* locus (*Osr1*^*LacZ*^, Fig. S1A), which allows for enzymatic signal amplification. The expression of the *Osr1*^*LacZ*^ allele recapitulated the developmental expression pattern of *Osr1* (Fig. S1B). Using the Osr1^LacZ^ line, we observed the presence of a low number of Osr1^+^ cells in the interstitium of several muscles examined (Fig. 1A). We noted that these cells are also positive for PDGFRα (Fig. 1B). FACS cytospin of FAPs (lin^−^;α7-integrin^−^;Sca1^+^) isolated from whole hindlimb muscle of *Osr1*^*LacZ*^ micerevealed that approx. 4.5% of adult FAPs were Osr1-β-Gal^+^ (Fig. 1C, Fig. S2). No β-Gal Signal wasdetected in lin^−^;α7-integrin^−^;Sca1^−^ cells (double negative, DN cells) or lin^−^;α7-integrin^+^;Sca1^−^ cells (satellite cells, SC) (Fig. 1C). We complemented this approach by isolating FAPs from *PWl*^*LacZ*^ animals (lin^−^;PW1^+^;PDGFRα^+^); this protocol yielded a population that completely overlapped FAPs (Pannerec et al., 2013). Low abundance of *Osr1* mRNA in adult FAPs was confirmed by semiquantitative PCR (Fig. 1D). This suggests that *Osr1* is expressed in a small proportion of adult FAPs and is consistent with deep RNA sequencing data from adult resident FAPs showing low *Osr1* mRNA expression (Ollitrault et al. in preparation).

**Figure 1.**
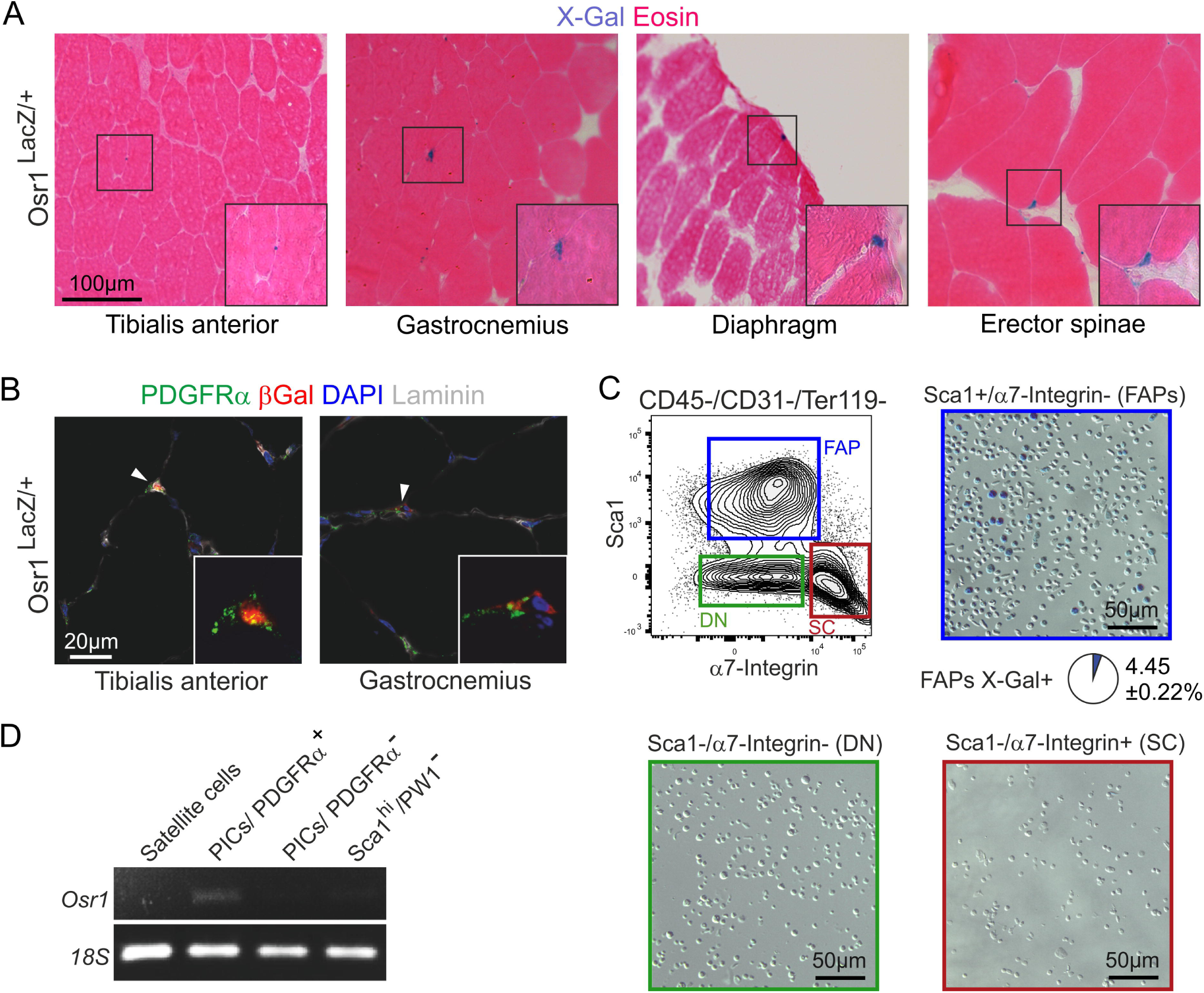
Osr1 is sporadically expressed in adult FAPs. **(A)** X-Gal staining on *Osr1*^*LacZ/*^+^^ several adult mouse muscles muscle shows sporadic X-Gal^+^ cells. **(B)** X-Gal positive cells also express PDGFRα. **(C)** Isolation of CD45^−^;CD31^−^;Ter119^−^;α7integrin^−^;Sca1^+^ FAPs from adult uninjured tibialis anterior of Osr1^LacZ^ mice followed by cytospin shows that only FAPs express Osr1 (n=3 animals). **(D)** Semiquantitative PCR on interstitial populations isolated from adult PW1^LacZ^ mice. Faint Osr1 expression can be detected in PDGFRα^+^ PICs.

### Osr1 expression is induced to high levels upon acute injury

Freeze-pierce injury performed on *Osr1*^*LacZ*^ tibialis anterior (TA) muscle led to an accumulation of Osr1^+^ cells in the injured region 3 and 5 days post injury (dpi) (Fig. 2A). We used both *Osr1*^*GCE*^ and *Osr1*^*LacZ*^ alleles to analyze which cells initiated *Osr1* expression. First, FAPs were FACS isolated from *Osr1*^*GCE*^ mice as lin^−^;α7-integrin^−^;Scal^+^ (Joe et al., 2010; Fig. S3A, B) and analyzed for Osr1-eGFP expression. Using this allele, we found that in uninjured muscle approx. 3.3% of FAPs expressed Osr1-eGFP (Fig. 2B), in agreement with results obtained from the *Osr1*^*LacZ*^ line (Fig. 1C). The numbers of Osr1-eGFP^+^ FAPs increased upon injury to 17 – 19% (approx. 5 – 6 fold increase) at 3, 5 and 7 dpi (Fig. 2B). At 10 dpi, the fraction of Osr1-eGFP^+^ FAPs decreased (8,5%; Fig. 2B). We note that isolation of FAPs from the whole TA muscle yields FAPs from non-injured and injured regions, whereas the concentration of activated FAPs in the injured region is higher. Osr1-eGFP was exclusively expressed in lin^−^;α7-integrin^−^;Scal^+^ FAPs and not detected in lin^−^;α7-integrin^+^;Scal^−^ SCs or Iin^−^;α7-integrin^−^;Sca1^−^ DN cells at 5 dpi (Fig. S3C). Cytospin analysis of all lin^−^;α7-integrin^−^;Sca1^+^ FAPs (i.e. GFP^+^ and GFP^−^) followed by immunolabeling for PDGFRα^+^ confirmed that after injury 18 – 20% of PDGFRa^+^ FAPs were Osr1-GFP^+^ at 3, 5 and 7 dpi, while this ratio declined to 8,5% at 10 dpi (Fig. 2C). In contrast, Osr1^+^ FAPs were almost completely positive for PDGFRα at all time points analyzed (Fig. 2C, Fig. S4A). Both the Osr1^+^ and the Osr1^−^ fraction of Scal^+^ FAPs overlapped partly with Tcf4 expression (Fig. 2C, Fig. S4A). We note that adult resting FAPs highly overlapped with the Tcf4^+^ population (Murphy et al., 2011; Vallecillo-García et al., 2017), while this was not the case after injury in our analysis. Moreover, the expression of Tcf4 did not correlate with Osr1 expression in FAPs after acute injury, suggesting dynamic changes in the FAP population during regeneration. We further noted expression of Tcf4 in α7-integrin^+^ myogenic cells (Fig. S4B), in agreement with Tcf4 expression in a fraction of developmental myoblasts (Mathew et al., 2011).

**Figure 2.**
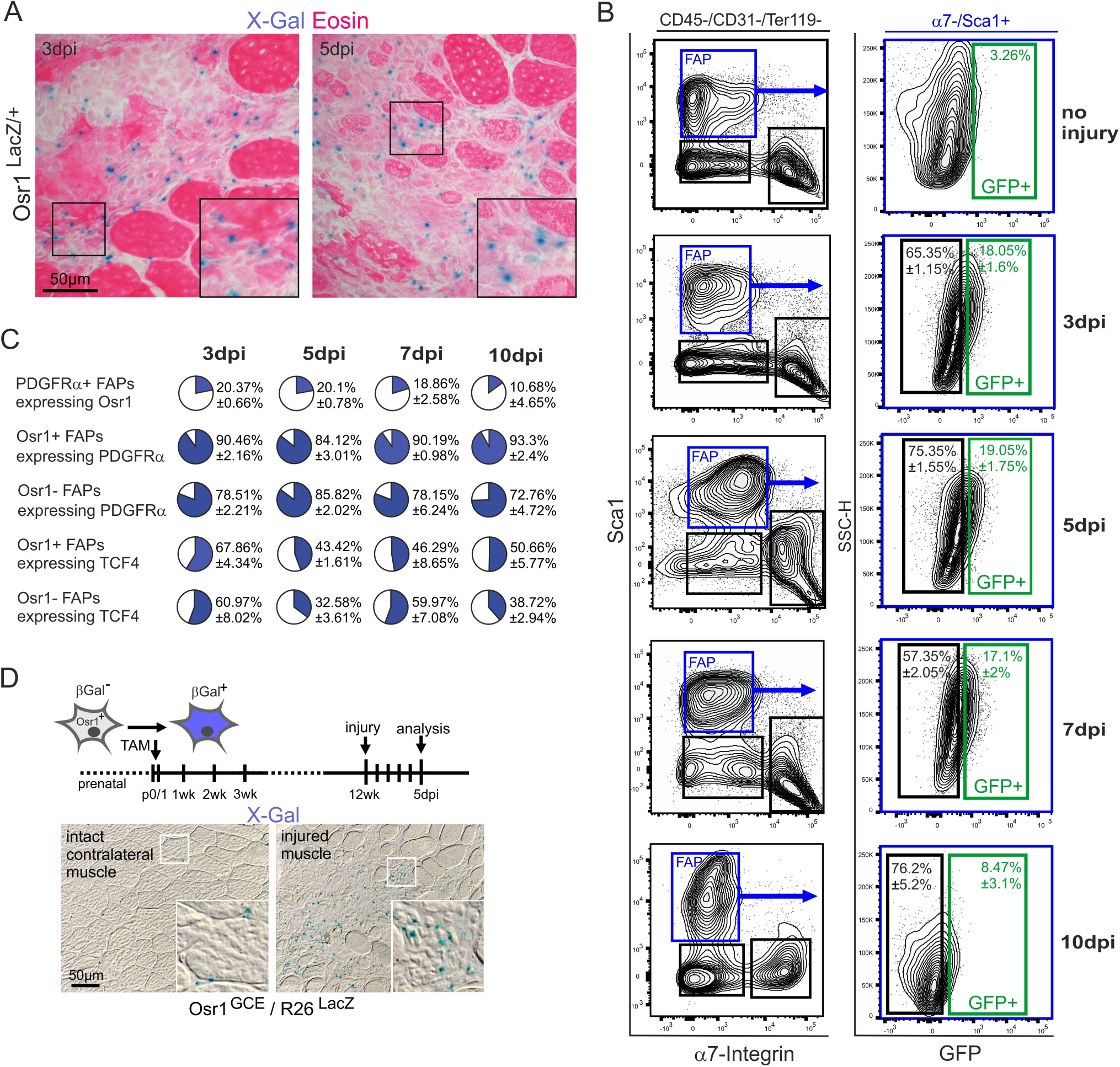
Osr1 expression is reactivated in FAPs upon acute injury. **(A)** X-Gal staining on *Osr1*^*LacZ/*^+^^ adult tibialis anterior muscle 3 and 5 days post injury (dpi) shows accumulation of *Osr1*-LacZ positive cells in the injury region. **(B)** FACS analysis of FAPs isolated at 3, 5, 7 and 10 dpi from Osr1^GCE/+^ mice for Osr1-GFP expression (n=3 animals for each time point). **(C)** Quantification of cytospin analysis of GFP^+^ and GFP^−^ FAPs isolated at indicated time points after injury from Osr1^GCE/^+^^ mice (n=3 animals). **(D)** Long-term lineage tracing using Osr1^GCE^;R26^LacZ^ animals Tamoxifen pulsed at p0 and p1 shows that the progeny of Osr1^+^ developmental cells expand in the injury region upon acute muscle injury in adult animals.

To corroborate these findings, FAPs were isolated from *Osr1*^*LacZ*^ animals as lin^−^;Scal^+^;CD34^+^ and analyzed via FACS for LacZ and PDGFRα expression (Fig. S5). Since LacZ staining can generate high background, we gated conservatively likely leading to exclusion of positive cells (Fig. S5B). Regardless, we could confirm *Osr1*-LacZ expression in 3 dpi lin^−^;Scal^+^ FAPs, and that LacZ^+^;Scal^+^ FAPs strongly overlapped with PDGFRα expression (Fig. S5C). Adult FAPs originate from a developmental Osr1^+^ lineage (Vallecillo-García et al., 2017). Consequently, progeny of developmental Osr1^+^ cells expanded in the injury region upon acute injury (Fig. 2D). Taken together, these data show that Osr1 expression is induced upon muscle injury specifically in FAPs within the injured region.

### Osr1 expression identifies injury-activated FAPs that contribute to adipogénie infiltration and postinjury resident FAPs

An initial rapid induction of proliferation is a hallmark of FAP activation in response to injury (Joe et al., 2010; Lemos et al., 2015; Uezumi et al., 2010) which is followed by apoptosis (Lemos et al., 2015). Following injury, we noted Ki67-stained Osr1^+^ cells in tissue sections from *Osr1*^*LacZ*^ animals (Fig. 3A). Next, FAPs were isolated (lin^−^;α7-integrin^−^;Sca1^+^) from Osr1^*GCE*^ animals at 3, 5 and 7 dpi. Osr1^+^ FAPs showed a significantly higher fraction of Ki67^+^ cells than Osr1^+^ FAPs (Fig. 3B). In addition, apoptotic cells detected by immunolabeling for cleaved caspase 3 were exclusively found in the Osr1^+^ FAP population at 7 dpi (Fig. 3C) consistent with the proposal that Osr1^+^ cells are activated in response to injury to undergo cell cycle entry as well as apoptosis.

**Figure 3.**
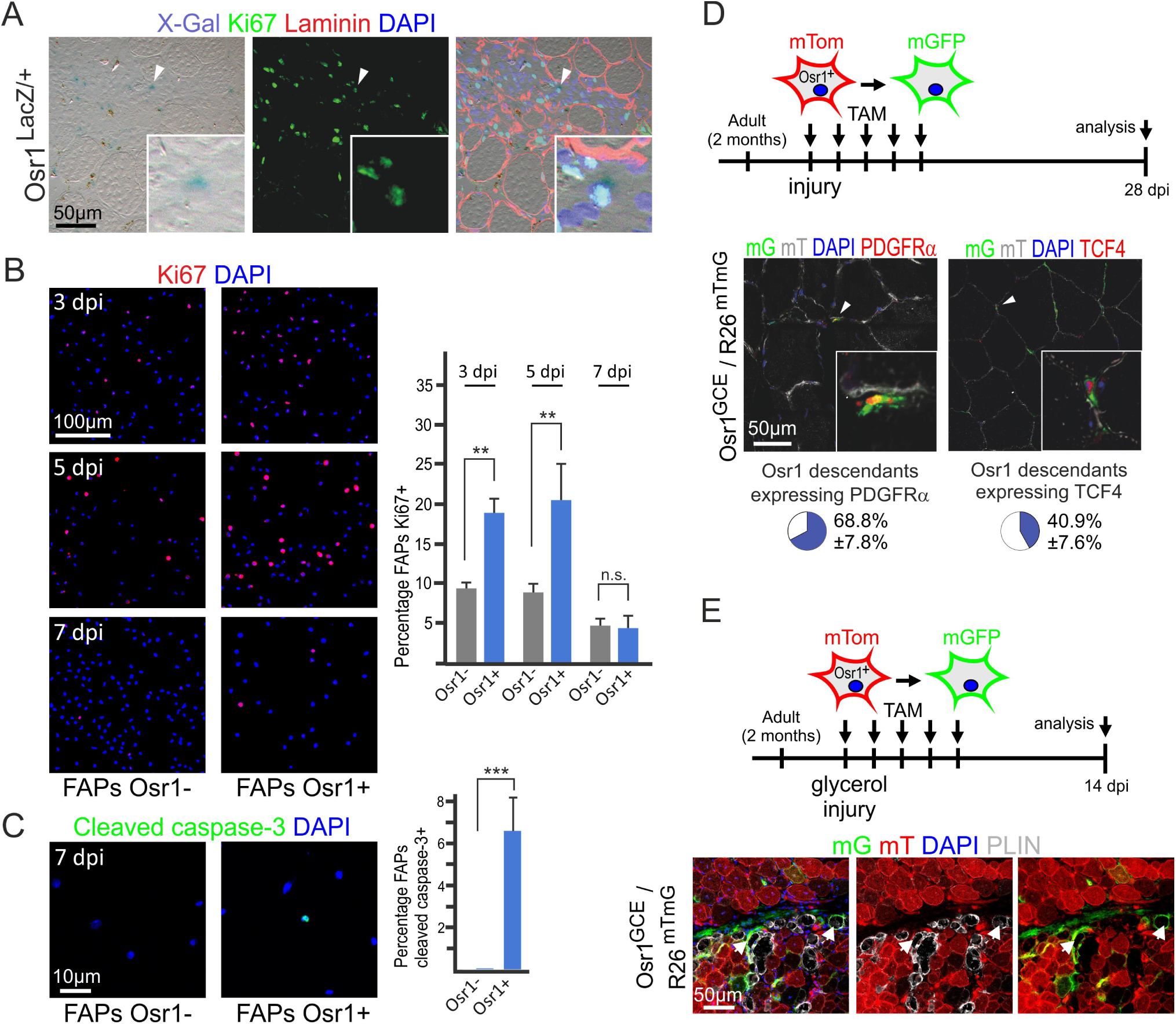
Osr1 marks injury-activated fibro-adipogenic progenitors (FAPs) **(A)** X-Gal^+^ (*Osr1* expressing) cells in the injury region (Osr1^LacZ/+^ animal; 3dpi) are positive for the cell cycle marker Ki67. **(B, C)** Cytospin of lin^−^;α7-integrin^−^;Sca1^+^ FAPs FACS-isolated at indicated days post injury (dpi) and separated into GFP^+^ and GFP" populations were stained for the proliferation marker Ki67 **(B)** and the apoptosis marker cleaved caspase 3 **(C)** (n=3 independent experiments). Quantification shown right. **(D)** Lineage tracing of Osr1^+^ cells in Osr1^GCE^;R26^mTmG^ animals Tamoxifen-pulsed for five consecutive days after freeze-pierce injury. Contribution of Osr1^+^ cells to PDGFRa^+^ and Tcf4^+^ interstitial cells was analyzed 28 days post injury (dpi). Quantification is shown below (n=3 animals). **(E)** Lineage tracing of Osr1^+^ cells in Osr1^GCE^;R26^mTmG^ animals Tamoxifen-pulsed for five consecutive days after glycerol injection injury. Contribution of Osr1^+^ cells to ectopic adipose tissue labelled for Perilipin (PLIN) was analyzed at 14 dpi. Data are represented as means +/-SEM; t-test: * = P<0,05, ** = p<0,01, *** = p<0,005, n.s. = not significant.

We next tested the suitability of the Osr1^GCE^ allele to trace the fate of activated FAPs. We genetically labelled Osr1^+^ cells in *Osr1*^*GCE*^;*R26*^*mTmG*^ mice for five consecutive days beginning with the day of injury (Fig. 3D). Since the Osr1^+^ FAP population expands in the injured region during this period, we would anticipate labeling of the activated FAP pool as compared to FAPs in uninjured muscle. Consistent with this prediction, we observed that pulsing Osr1^+^ cells for five days before injury resulted in low levels of labeling as compared to labeling post injury (Fig. S6). This also suggests that Osr1^+^ cells in adult uninjured muscle do not represent a specific subpopulation prone to quick expansion upon injury, rather, we propose that sporadic Osr1 expression in uninjured adult muscle results from activated FAPs engaged in focal repair, however this requires further investigation. Lineage tracing in *Osr1*^*GCE*^;*R26*^*mTmG*^ mice induced after injury was performed at 28 days after injury, where regeneration is almost completed, however the regenerated tissue can be recognized by centrally located myonuclei. The majority of Osr1^+^ cell progeny after injury was traced to interstitial PDGFRα^+^ cells representing resident FAPs (Fig. 3D). Osr1^+^ cells also gave rise to interstitial Tcf4^+^ cells (Fig. 3D). This suggests that Osr1^+^ activated FAPs return to the resident FAP pool after regeneration as well as to the Tcf4^+^ muscle connective tissue fibroblasts consistent with the proposal that FAPs are a primary source for fibrosis in degenerative disease (Contreras et al., 2016; Lemos et al., 2015; Mueller et al., 2016; Uezumi et al., 2011).

In addition to being a source of fibrotic tissue in pathologically remodeled skeletal muscle, FAPs are also proposed to be a source of fatty infiltration (Lemos et al., 2012; Uezumi et al., 2010; Uezumi et al., 2011) although this has not been conclusively demonstrated in situ. Therefore, we tested the adipogénie potential of Osr1^+^ injury-activated FAPs, by glycerol injury in *Osr1*^*GCE*^;*R26*^*mTmG*^ mice, which results in fatty infiltration (Pisani et al., 2010) (Fig. 3E). 14 days after injury, the regenerating region contains infiltrating adipocytes which are mG positive (Fig. 3E). Taken together, these results provide a key confirmation for endogenous FAP fibrotic and adipogénie fates *in situ*.

### Osr1 marks a transient population of juvenile Scal^+^ cells

We noted previously that Osr1 expression fades in early postnatal life (Vallecillo-García et al., 2017), which is a period of still active myogenesis. Furthermore, early postnatal development represents a dynamic phase in which the transition of developmental progenitors to resident stem cells is accomplished. On this background, we finally re-analyzed Osr1 expression in juvenile mice. In young mice (p7) β–Gal from the *Osr1*^*LacZ*^ allele was expressed in muscle interstitial cells (Fig. 4A), most of a interstitial PDGFRα^+^ cells that likely are FAPs / FAP progenitors and hence should also express Seal (Pannerec et al., 2013; Uezumi et al., 2010).

**Figure 4.**
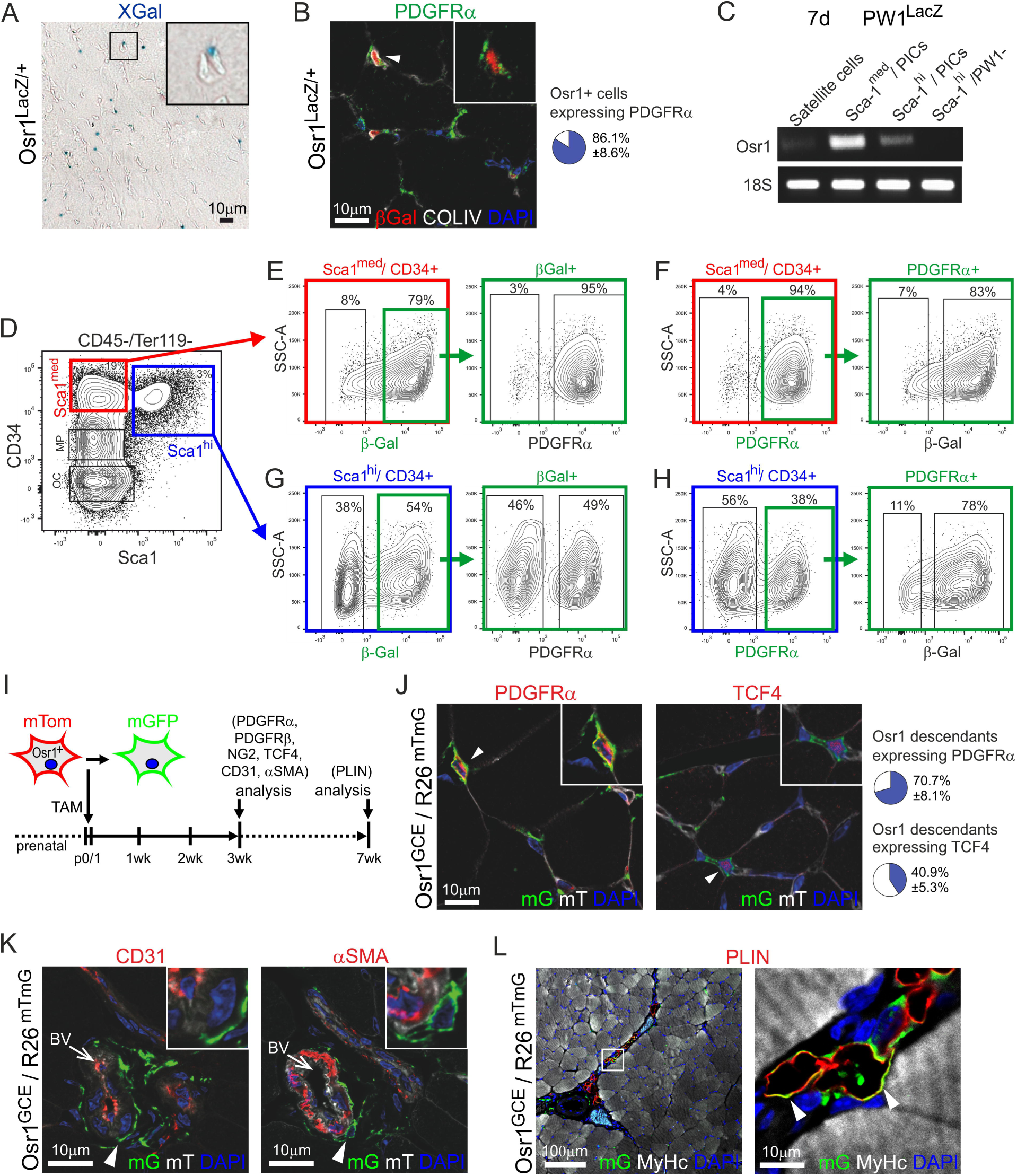
In juvenile mice Osr1 is expressed in a transient Scal^med^ population. **(A)** X-Gal staining on 7 day old *Osr1*^*LacZ/*^+^^ tibialis anterior muscle shows Osr1-LacZ^+^ interstitial cells. **(B)** Osr1-LacZ positive cells are mostly positive for PDGFRα in situ (n=3 animals). **(C)** Semiquantitative PCR on interstitial populations isolated from 7 day old PWl^LacZ^ mice. Strong *Osr1* expression can be detected in Scal^med^ PICs, while faint *Osr1* expression can be detected in Sca1^hi^ PICs. (D) Isolation of CD34^+^/Scal^+^ cells from 7 day old tibialis anterior muscles; the Scal^med^ and Sca1^hi^ populations are CD34^+^/Sca1^+^ cells from 7 day old tibialis anterior muscles; the Sca1^med^ and Sca1^hi^ populations are discernible. **(E, F)** Analysis of the Sca1^med^ population. **(E)** Sca1^med^ cells are mostly PDGFRα positive; Scal^med^ PDGFRa^+^ cells mostly express β-Gal. **(F)** Scal^med^β-Gal^+^ cells express PDGFRa. **(G, H)** Analysis of Sca1^hi^ cells. **(G)** Sca1^hi^ cells are only in part positive for PDGFRα; Sca1^hi^;PDGFRα^+^ cells mostly express β-Gal. **(H)** Only approx. half of Sca1^h1^ cells express β-Gal; Sca1^h1^β-Gal^+^ cells are also only in part PDGFRα^+^. **(I-L)** Lineage analysis of perinatal Osr1^+^ cells. **(I)** Lineage tracing strategy. Tamoxifen was administered at p0/p1, analysis was performed at 3 weeks of age; adipogénie differentiation was analyzed at 7 weeks of age. **(J)** Osr1^+^ progeny majorly express PDGFRα, but in part also TCF4. Quantification shown right (n= 2 animals). **(K)** Osr1^+^ cells do not give rise to CD31^+^ endothelial cells or αSMA^+^ smooth muscle cells. **(L)** Osr1^+^ cells give rise to muscle interstitial adipocytes labelled for Perilipin (PLIN). Data are represented as means +/-SEM

Juvenile Scal^+^ cells had been analyzed on the background of muscle interstitial PICs (Pannerec et al., 2013). This study showed that juvenile Scal^+^ cells can be subdivided into two populations: one with low/medium Sca1 expression levels (Seal^med^) and a population with high Seal expression (Scal^hi^) that persists throughout life and completely overlaps with FAPs. We therefore FACS-isolated Sca1^med^;PW1^+^ and Sca1^hi^PW1^+^ PICs from 7 day old *PW1*^*LacZ*^ mice as previously described (Pannerec et al., 2013) and analyzed *Osr1* expression by semiquantitative PCR. Strong *Osr1* expression was seen in Scal^med^, whereas lower levels of expression were detected in Scal^hi^ PICs. No detectable levels of expression were found in Sca1^hi^PW1^+^ cells or in satellite cells (Fig. 4C)

To further characterize Osr1^+^ juvenile cells, we first FACS isolated lin^−^;CD34^+^;Scal^med^ and lin^−^;CD34^+^;Sca1^hi^ cells from *Osr1*^*LacZ*^ mice (Fig. 4D). The majority of Sca1^med^ cells were β-Gal positive (79%), of which almost all Sca1^med^;β-Gal^+^ cells were PDGFRa positive (95%; Fig. 4E). In contrast, Sca1^med^;PDGFRα^+^ cells were mostly β-Gal positive (94%; Fig. 4F). The Sca1^hi^ population contained distinct and separable β-Gal^+^ and β-Gal^−^ subpopulations. Interestingly, of the Sca1^hi^;β-Gal^+^ population only approx. 50% expressed PDGFRα (Fig. 4G). In contrast, most of the Sca1^hi^;PDGFRα^+^ cells were β-Gal^+^ (78%; Fig. 4H), showing a prevalence for Osr1 expression in the PDGFRα fraction. Taken together, these results show that Osr1 is expressed primarily in interstitial Sca1^med^;PDGFRα^+^ cells and to lower extent in Sca1^hi^;PDGFRα^+^ cells in juvenile muscle.

We next re-analyzed the developmental fate of perinatal Osr1^+^ cells by genetically labeling the Osr1 lineage in *Osr1*^*GCE*^*;R2*^*mTmG*^ mice. Administration of Tamoxifen (TAM) at p0 and p1 was used to developmentally trace Osr1^+^ cells (Fig. 1I). At day p21, Osr1^+^ progenitors contributed to the muscle interstitial PDGFRα cell pool (approx. 70%; Fig. 1J) in line with our previous demonstration that adult FAPs derive from these developmental Osr1^+^ cells (Vallecillo-García et al., 2017). Osr1 descendants were also present in cells expressing the muscle connective tissue fibroblast marker, TCF4, although at a markedly lower number (Fig. 1J). No contribution of Osr1^+^ progenitors was seen in αSMA^+^ vascular smooth muscle or CD31^+^ endothelium (Fig. 1K). In addition, Osr1^+^ progenitors gave rise to muscle interstitial adipocytes (Fig. 1L). This shows that juvenile Osr1^+^ cells maintain plasticity and that interstitial fibroblasts as well as interstitial adipocytes derive from this pool in addition to FAPs. During embryonic development, Osr1 is expressed in Seal^−^ cells whereas Seal-expressing muscle interstitial cells arise during late fetal development, concomitant with the appearance of Scal^+^/Osr1^+^ cells (Vallecillo-García et al., 2017). While it remains unresolved whether Osr1^+^ cells acquire Seal expression during fetal development or whether the Scal^+^ cells arise *de novo*, our observation that Osr1 is predominantly expressed in a transient population of Scal^med^ FAPs during postnatal development has interesting implications. Specifically, we note that Osr1 is expressed in muscle interstitial mesenchymal cells throughout embryonic and postnatal myogenesis, first in Seal^−^, later in Scal^med^ and to a lesser degree in Sca1^hi^ cells. This pattern of expression suggests a lineage continuum in which Sca1^hi^ cells are derived from Scal^med^ cells, and that downregulation of Osr1 expression is required for this differentiation step. Interestingly, both the disappearance of Scal^med^ cells and the downregulation of Osr1 expression correlate with the termination of postnatal muscle growth driven by incorporation of new nuclei into myofibers (Pannerec et al., 2013; White et al., 2010). These data together with the observation of re-activation of Osr1 expression in injury-activated FAPs leads to the proposal that Osr1 expression marks FAPs or their progenitors during periods of active myogenesis. During embryonic myogenesis, Osr1^+^ cells provide a pro-myogenic niche for myogenic progenitors that promotes myogenic cell proliferation and survival. (Vallecillo-García et al., 2017). Similarly, adult FAPs promote myogenesis *in vitro* (Joe et al., 2010) and Tcf4^+^ cells, which in part overlap with FAPs, are required for efficient muscle regeneration (Murphy et al., 2011). The re-activation of Osr1 expression further suggests that adult FAPs reactivate a developmental program to support tissue regeneration.

In summary, we show that Osr1 is the first specific marker identified for injury activated FAPs. Given the key role for FAPs in promoting proper muscle regeneration, the ability to lineage track these cells will be invaluable for designing approaches to optimize muscle repair and target the muscle stem cell niche.

## Materials and methods

### Mice

Mice were maintained in an enclosed, pathogen-free facility. The targeting construct for the *Osr1* multifunctional allele was electroporated into G4 mouse ES cells (George et al., 2007). The transgenic locus was confirmed by Southern blotting and Sanger sequencing. Successfully recombined clones were subjected to tetraploid aggregation. Mice were crossed back to wild type C57BL/6j mice for at least 6 generations before establishing the line. The following mouse lines were described before: Osr1^GCE^ (Mugford et al., 2008); R26^LacZ^ (Soriano, 1999); R26^mTmG^ (Muzumdar et al., 2007). Mouse experiments were performed in accordance with European Union regulations (Directive 2010/63/EU) and under permission from the Landesamt für Gesundheit und Soziales (LaGeSo) Berlin, Germany (Permission numbers ZH120, G0240/11, G0114/14, G0209/15, G0268/16).

### Cell lineage tracing

Lineage tracing of neonatal Osr1^+^ cells was performed in Osr1^GCE/^+^^;R26^lacZ/^+^^ neonates by subcutaneous injection of Tamoxifen (Sigma Aldrich; solved in 90 % (v/v) sunflower oil/ 10 % (v/v) ethanol) into the neck fold (75 μg/ g body weight). Lineage tracing of injury-activated Osr1^+^ FAPs was performed in Osr1^GCE/^+^^;R26^mTmG/^+^^ or Osr1^GCE^;R26^lacZ^ mice by intraperitoneal injection of Tamoxifen at the day of injury and the next 4 following days (3 mg per injection time point). For lineage tracing of Osr1^+^ cells before injury, *Osr1*^*GCE/^+^*^*;R26R*^*LacZ/+*^ and *Osr1*^*GCE/+*^*;R26R*^*mTmG/+*^ animals were injected with 3mg Tamoxifen for 5 consecutive days one week before injury.

### Muscle injury

Injury was applied to the tibialis anterior muscle of 3-5 months old Osr1^GCE^, Osr1^lacZ^, Osr1^GCE^;R26^mTmG^ or Osr1^GCE^;R26^lacZ^ mice using the freeze-pierce technique. Mice were anesthetized by intraperitoneal injection of 10 % (v/v) ketamine / 2 % (v/v) xylazine (Rompun^®^ 2%) in sterile PBS (5 μl / g body weight). Mice were kept on a heating plate warmed to 37°C throughout the procedure. The skin above the Tibialis anterior muscle was opened and the muscle was longitudinally pierced 5 times using a 0.7 mm liquid nitrogen cooled syringe needle.

For glycerol injury mice were anesthetized as described above. Skin above the Tibialis anterior muscle was opened and 25μΙ of 50% v/v glycerol/sterile PBS were injected into the muscle.

### Histology, antibody labeling

Muscle was dissected, immediately embedded in 6 % (w/v) gum tragacanth (Sigma Aldrich) dissolved in H_2_0 and snap frozen in liquid nitrogen cooled isopentane (-160 °C). Frozen tissue was sectioned at 7 μm and fixed using 4 % (w/v) PFA in PBS for 5 min at RT.

Permeabilization of sections was performed in 0.3% (v/v) Triton ☓-100 (Sigma Aldrich) in phosphate buffer (PBS) for 10 min. Sections from adult tissues were blocked with 5% (w/v) bovine serum albumin (Sigma Aldrich) in PBS. Sections then preincubated with 5% horse serum, 5mg ml/^1^ blocking reagent (Perkin Elmer) and 0.1% Triton ☓-100 in PBS for 1h at RT. Primary antibodies were applied in the same solution and incubated at 4 °C overnight, followed by secondary antibody staining for 1h at room temperature. Antibodies are listed below. Counterstaining was performed with 5μg μL^-1^ 4',6-diamidino-2-phenylindole (DAPI; Invitrogen), slides were mounted with FluoromountG (SouthernBiotech). Antibodies are listed in Additional file 1, Supplementary tables 1 and 2.

For beta galactosidase detection, sections or cells were fixed as above and incubated at 37°C overnight in X-Gal (0.16 % (w/v) X-Gal, 5mM K_3_Fe(CN)_6_, 5 mM K_4_Fe(CN)_6_, 2 mM MgCl_2_ in PBS). Beta galactosidase development was stopped by washing in PBS. For whole-mount X-Gal stainings embryos were fixed in X-Gal fixing solution (1 % (w/v) Formaldehyde, 0.2 % (w/v) Glutaraldehyde, 0.02 % (v/v) NP-40, 1 % (v/v) PBS in bidest H_2_O) for 1 h at 4°C, then stained for 24 h at 37°C in X-Gal staining solution.

### Cell isolation and flow cytometry

Cell isolation and labeling was essentially performed as described in (Vallecillo-García et al., 2017). Isolation of cells from Osr1^GCE/^+^^ mice was performed as follows: Briefly, whole hind limb or *tibialis anterior* muscles were carefully isolated, roughly minced and digested in high-glucose DMEM medium containing 10% fetal calf serum (FCS, Biochrom), 1% Penicillin Streptomycin solution (P/S; 10000 U/ml) and 2,5 mg/ml Collagenase A (Roche) for 75 min at 37°C with vigorous shaking. 2 IU/ml of Dispase II (Sigma Aldrich) were added to the digestion solution and muscle lysates were digested for further 30 min. Muscle slurries were passed 10 times through a 20G syringe (BD Bioscience) and a 70-μm cell strainer. Cells were collected by centrifugation at 300g for 5 min and resuspended in staining buffer consisting of 500μΙ Hank′s balanced salt solution (HBSS, Thermo Fisher scientific), 0.4 % bovine serum albumin (Sigma Aldrich) and 20μg/ml Gentamycin (Serva Electrophoresis). Cells were stained on ice for 30 min and washed twice with staining buffer previous to FACS sorting. Propidium iodide was used as a viability dye.

Isolation of cells from Osr1^LacZ/+^ animals was performed as follows: freshly dissected muscle tissue was minced and digested in HBSS (Gibco) supplemented with 2.4 U/ml Dispase || (Roche), 2 μg/ml Collagenase A (Roche), 0.4 mM CaCl_2_, 5 mM MgCl_2_, 10 ng/ml DNase | (Roche) for 2 h at 37°C under agitation. Single cell suspension was obtained after 3 successive cycles of straining and washing in Washing Buffer consisting of HBSS containing 0.2% (w/v) BSA (Sigma Aldrich), 1% (v/v) penicillin-streptomycin, 10 ng/ ml DNAse I and 10% (v/v) mouse serum. Cells were incubated with primary antibodies for 1 h on ice. The suspension was subjected to 2 washing steps, resuspended in Washing Buffer and to LacZ reporter staining using C_12_FDG (Life Technologies). For lacZ staining C_12_FDG was added to the cell suspension to a final concentration of 600 μΜ and incubated for 30 min at 37°C. 2 washing steps followed before proceeding to FACS analysis.

Cells purified by sorting were cytospun to coverslips. Coverslips were coated with poly-L-lysine by incubation with a 10-fold solution of poly-lysine in bidest H_2_0 for 1 h at RT, rinsed twice in bidest H_2_O and air dried. Purified cells were added to prepared slides and allowed to adhere for 1 h at 4°C. Supernatant was removed by centrifugation at 50 rcf for 5 min at 4°C, cells were fixed using 4% (w/v) PFA in PBS for 5 min at RT and permeabilized with 0.3% (v/v) Triton X-100 in PBS for 5 min at RT. Antibodies are listed in Additional file 1, Supplementary tables 1 and 2.

Sorts and analyses were performed on a FACS Aria II (BD Biosciences). Sorting gates were defined based on unstained controls. Data were collected using FACSDIVA software. Biexponential analyses were performed using FlowJo 10 (FlowJo LLC) software. Analysis was performed on three independent biological replicates.

### Cell quantification

Quantification of FACS-isolated Osr1^+^ cells after immunolabelling (cytospin) was performed on at least three independent biological replicates (i.e. cells FACS isolated from three different animals). Quantifications of cytospun cells were made from two areas of 0.81 mm^2^ per replicate.

Student's t-test was performed using Prism 5 (GraphPad) software. Error bars in all figures, including supplementary information, represent the mean ± standard error of the mean (s.e.m).

### Microscopy

Images were acquired using a Zeiss LSM700 confocal microscope, a Leica DMR or Leica DMi8 microscope. Bright field images of whole-mount embryos were taken with a Leica Leica MZ12 stereo microscope. Images were captured using Axio Vision Rel. 4.8 and Zen 2010 (Zeiss) or LAS X (Leica).

## Acknowledgements

We thank the animal facility of the Max Planck Institute for Molecular Genetics, Berlin for expert support, especially Karol Macura, Judith Fiedler, Lars Wittler, Andrea König, Katja Zill and Ludger Hartmann.

## Author’s contributions

DAS and SS conceived and designed the study. JS, PVG, SvHS and DO performed experiments and collected data. JS, PVG, SvHS, DO, GM, DAS and SS performed data analysis and interpretation. ANE generated the Osr1-LacZ knock-in construct and HS generated the Osr1-LacZ knock-in ES cells. DAS and SS wrote the manuscript with help from JS and GM.

